# Mesenchymal predominance in olfactory epithelium–derived cultures limits modeling of neurodevelopmental brain disorders

**DOI:** 10.64898/2026.04.23.720382

**Authors:** Xoel Mato-Blanco, Silvia Beltramone, Marta Barrera-Conde, Emma Veza-Estevez, Zenaida Piñeiro, Ares Ramos, Anna Mané, Arnau Cendón, Maria José Algora, Mayte Gomáriz, Claudia Sánchez-Aldabó, Amira Trabsa, Vanessa Sanchez-Gistau, Pilar Àlvarez, Rafael de la Torre, Gerard Muntané, Patricia Robledo, Gabriel Santpere

**Affiliations:** Hospital del Mar Research Institute, Parc de Recerca Biomèdica de Barcelona (PRBB), 08003 Barcelona, Catalonia, Spain; Institut de Salut Mental, Hospital del Mar, Psychiatry Department, 08003 Barcelona, Spain; Hospital Universitari Institut Pere Mata, Reus, Catalonia, Spain; Institut de Recerca Biomèdica Catalunya Sud (IRBCatSud), Reus, Catalonia, Spain Universitat Rovira i Virgili, Reus, Catalonia, Spain; Centro de Investigación Biomédica en Red de Salud Mental (CIBERSAM), Instituto de Salud Carlos III, Madrid, Spain; Institut de Biologia Evolutiva (UPF-CSIC), Department of Medicine and Life Sciences, Universitat Pompeu Fabra, Parc de Recerca Biomèdica de Barcelona, Barcelona, Catalonia, Spain; Department of Neuroscience, Yale School of Medicine, New Haven, CT, 06510, USA

**Author notes:** Authors equally contributed (Xoel Mato-Blanco, Silvia Beltramone).

**Keywords:** Olfactory neuroepithelium, horizontal basal cells, globose basal cells, neurogenesis, mesenchymal cells, brain disorders

## Abstract

The human olfactory epithelium (OE) represents a lifelong source of neural progenitor cells and has been proposed as an accessible model to investigate molecular alterations associated with neurodevelopmental disorders in postnatal individuals. Globose basal cells are considered the immediate neuronal progenitors within the OE, and several studies have attempted to culture these cells from nasal exfoliates. However, the actual contribution of neurogenic lineages in these cultures remains largely unquantified. Here, we cultured human nasal explants using an established protocol and characterized the resulting cell populations by immunohistochemistry and single-cell RNA sequencing. Integration with primary in vivo OE datasets revealed that these cultures are predominantly composed of mesenchymal-like cells, with limited representation of globose basal cells and neurons, and low expression of canonical neuronal markers. Using curated gene sets associated with neurodevelopmental disorders and malformations of cortical development, we assessed the extent to which disease-relevant transcriptional programs are captured in OE-derived cultures. While disease-associated genes are enriched in neurogenic lineages in vivo, their representation in mesenchymal cells is reduced. Together, our results challenge the assumption that standard OE culture systems faithfully model neurogenic compartments and suggest that current approaches may need refinement to recover neurogenic lineages.

## Introduction

The progeny of neural stem cells (NSC) constitute the neuronal and macroglial component of the entire nervous system. Disruption of NSC have been linked to multiple brain pathologies, from cortical malformations to neuropsychiatric disorders [1–5]. While genes associated to many brain disorders show enrichment in neuronal signatures [6, 7], a substantial fraction of disease-associated genes are also expressed in different types of NSCs and play disease-specific roles during neurogenesis, gliogenesis and progenitors’ maturation [7]. Genome-wide association studies, refined by functional genomics atlases, have linked loci associated to neuropsychiatric disorders, such as autism and schizophrenia, with fetal-specific regulatory elements and prenatal eQTLs [6, 8, 9], suggesting that the evaluation of early cell types during neurodevelopment might offer not only etiological insights, but also better diagnostic precision and power.

Because these developmental windows are inaccessible in humans, induced pluripotent stem cells (hIPSCs) derived from patients and controls have been widely used to interrogate early molecular phenotypes associated with pathology. However, the procedure has limitations, including variability in differentiation, inter-line heterogeneity, and restricted scalability for powered statistics. As the hIPSCs field continues to evolve, alternative human-derived NSC models have been explored. One such system is the olfactory epithelium (OE), a unique peripheral niche in which bona fide neurogenesis persists throughout the human lifespan. Owing to its ongoing cellular turnover and accessibility in living subjects, the OE provides a rare opportunity to investigate active neurodevelopmental and NSC-related processes postnatally. Indeed, OE-derived cultures have been used to capture disease-relevant molecular and signaling phenotypes as models of patient-specific peripheral cellular systems [10–22].

The OE is located on the nasal side of the lamina cribrosa, a perforated bone that supports the olfactory bulb and allows passage of olfactory axons, and continuously generates olfactory sensory neurons throughout life. These primary neurons are exposed to the external environment to detect odorants and relay signals to the brain, and are therefore subject to constant turnover [23]. Their cell bodies are organized beneath an apical layer of sustentacular and microvillar cells, which provide structural and secretory support, and are generated from basal progenitors residing in the basal lamina.

The basal compartment comprises two mitotically active populations: horizontal basal cells (HBCs) and globose basal cells (GBCs). Early studies from the 1970s and 1980s identified GBCs as the primary neuronal progenitors in the OE, while HBCs and sustentacular cells were considered distinct lineages [24–26]. However, subsequent work suggested broader lineage relationships, including the potential generation of sustentacular cells from HBCs [27]. HBCs have also been implicated in stem cell activity and epithelial regeneration, although they are generally considered a more quiescent population capable of giving rise to GBCs [28–30]. More recently, single-cell transcriptomic analyses of human OE have delineated a differentiation trajectory linking proneural GBCs to immature and mature olfactory neurons [31]. These neurogenic GBCs express canonical proneural basic helix–loop–helix transcription factors, including ASCL1, NEUROG1, and NEUROD1, consistent with earlier observations in mouse models [32–34]. Among these, NEUROD1 marks later stages of the lineage, being expressed at the transition from proliferating progenitors to early postmitotic neurons [34, 35].

Various approaches have been used to investigate the OE, typically obtained via nasal brushing of the olfactory cleft. Collected samples have been expanded in adherent conditions using DMEM-based media [10], embedded in extracellular matrix supports such as Matrigel [11, 12], or, more recently, cultured as neurospheres under FGF-enriched conditions [13, 36]. Neuronal-like cells have also been derived from these primary cultures using different differentiation protocols. Bulk homogenates from patient-and control-derived cultures have been analyzed using qPCR or bulk RNA sequencing to identify neurogenesis-related alterations associated with diverse conditions, including schizophrenia [11, 15, 16], bipolar disorder [18, 19], cannabis exposure [10, 20], and cystic fibrosis [15], among others. However, these studies generally assume that the relevant progenitor populations, namely HBCs and GBCs and their neuronal progeny, significantly contribute to the measured transcriptomes. In many cases, this assumption has not been directly validated.

Importantly, variability in sampling procedures and anatomical targeting of the OE (typically within the middle turbinate), even under endoscopic guidance, as well as differences in culturing conditions, can substantially influence the cellular composition of the collected material, thereby confounding downstream molecular analyses. Moreover, neuronal-like cells derived from these cultures may not necessarily recapitulate bona fide globose basal cell–driven neurogenesis in vivo, but instead reflect the differentiation potential of alternative mesenchymal or stromal populations present in the biopsy, as previously suggested [12, 37]. Consistent with this notion, transcriptomic analyses of neurosphere-derived cells have revealed a progressive shift toward stromal and smooth muscle–like signatures upon in vitro expansion [38].

Here, we focus on the GBC–to–neuron lineage within the human OE, with the aim of determining whether this neurogenic axis is preserved in in vitro culture systems derived from nasal exfoliates. We first perform a comprehensive cellular characterization of adherent OE cultures using immunostaining and single-cell RNA sequencing across samples obtained from multiple donors and independent laboratories. Leveraging signatures derived from primary human OE tissue, we then quantify the representation of GBCs and neuronal populations in vitro, and extend this analysis to previous bulk RNA-seq datasets from a total of 290 individuals using deconvolution approaches.

Having established the cellular composition of these cultures, we next assess the suitability of mesenchymal cells for modeling early pathogenic events associated with neurodevelopmental disorders. To this end, we use curated gene sets linked to malformations of cortical development (MCDs) and neurodevelopmental disorders (NDDs) [7], and examine their expression patterns across cell types in the human OE. By comparing the distribution and prevalence of disease-associated gene expression across neurogenic and non-neurogenic populations, we evaluate whether the cellular composition of OE-derived cultures is compatible with capturing neurogenesis-relevant molecular signals.

## Results

### Cellular characterization of ON derived cells in adherent culture

We obtained OE explants from six subjects (**Table S1**) using an established protocol [10]. To assess protocol reproducibility, we focused on samples obtained by three different operators, including two trained nurses and one otorhinolaryngologist from two independent centers (**Table S1**). Nasal exfoliates were cultured for three weeks and subsequently expanded through serial passaging. Cells were analyzed by fluorescent immunohistochemistry at passage 3 and, for a subset of two cultures, also at passage 5.

Cells were stained with TUJ1 antibody (class III β-tubulin, TUBB3), a tubulin isoform largely restricted to neurons and commonly used as a marker of neuronal lineage in cultures derived from nasal exfoliates [15, 17–21, 36]. In agreement with these previous reports, we observed widespread TUBB3 staining outlining the cellular cytoskeleton (**Figure 1A**). A large fraction of cultured cells also showed nuclear expression of KI67, indicating active proliferation.

**Figure 1.**
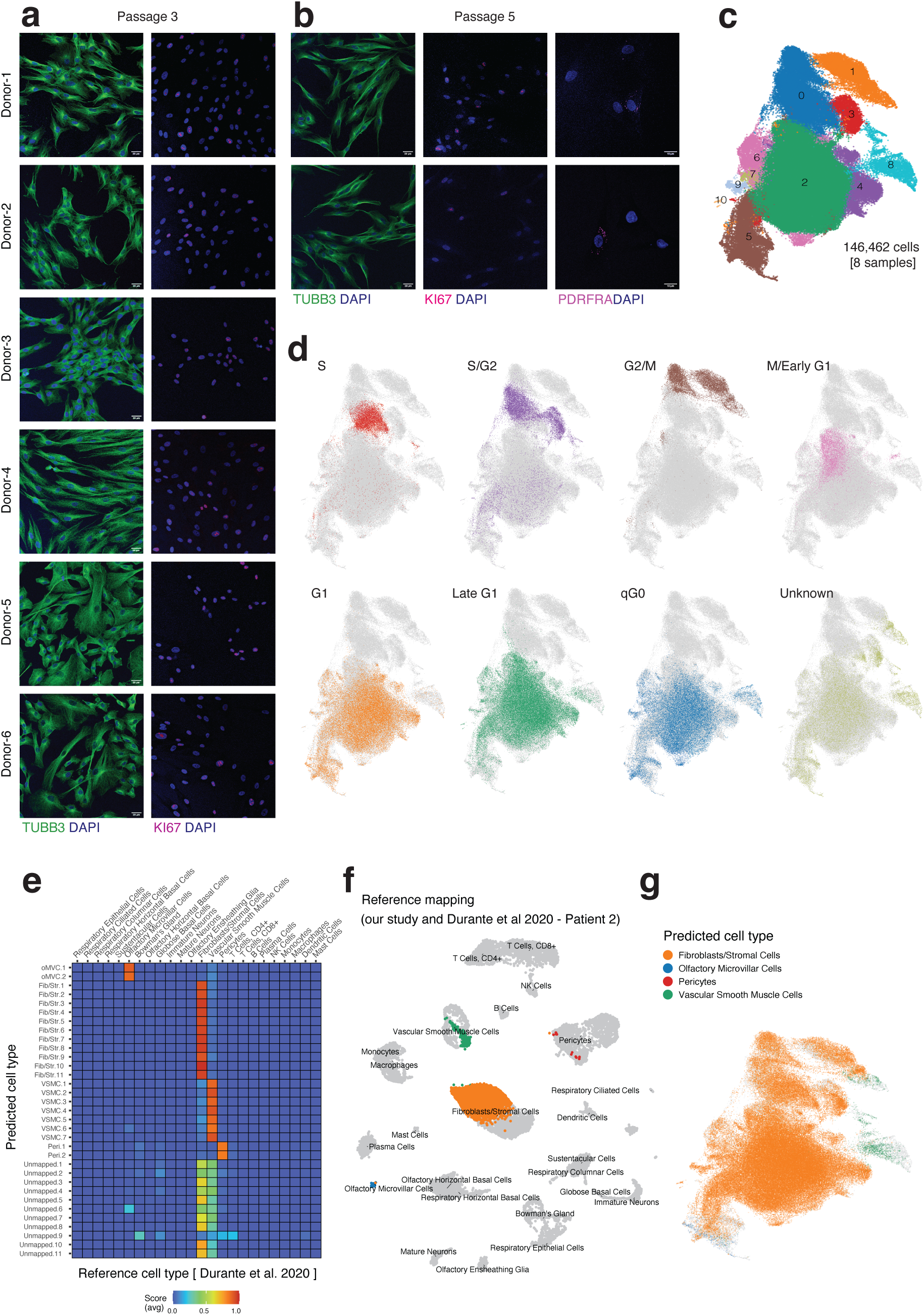
Adherent cultures derived from OE exfoliates display mesenchymal features and lack neurogenic populations. (**a**) Immunofluorescence staining of passage 3 cultures derived from nasal exfoliates of six individuals. Cells show widespread expression of TUBB3 (green) and nuclear staining with DAPI (blue). Despite TUBB3 positivity, cellular morphology is consistent with fibroblast-like cells. KI67 staining indicates that a large fraction of cells are actively proliferating. (**b**) Immunofluorescence staining at passage 5 showing persistent TUBB3 expression and proliferative activity (KI67), together with detection of PDGFRA (magenta), consistent with a mesenchymal/stromal identity. (**c**) UMAP representation of 146,462 single cells from eight samples after integration and high-resolution clustering, revealing 11 clusters. (**d**) Projection of cell cycle phase onto the UMAP, showing a continuum of cycling states (S, S/G2, G2/M, M/Early G1, G1, Late G1, qG0), indicating that cell cycle is a major source of transcriptional variation. (**e**) Heatmap showing label transfer scores between clusters identified in this study and reference cell types from the human olfactory OE [31], highlighting predominant alignment with fibroblast/stromal populations. (**f**) Reference mapping of the integrated dataset onto the human OE atlas, illustrating the correspondence between query cells and annotated in vivo cell types. (**g**) UMAP colored by predicted cell type following label transfer, showing that the majority of cells are classified as fibroblasts/stromal cells, with minor contributions from vascular smooth muscle cells, pericytes, and olfactory microvillar cells.

Despite the presence of TUBB3, cellular morphology was more consistent with fibroblast-like or mesenchymal cells, characterized by an extended soma, prominent filopodia and lamellipodia, and absence of polarity. In addition, the majority of cells appeared actively cycling. These observations suggest that, in this culture system, TUBB3 expression does not necessarily correspond to differentiated neurons but may also be present in proliferative mesenchymal-like cells. Consistent with this interpretation, a subset of cells at passage 5 was positive for PDGFRA, a marker associated with stromal and mesenchymal progenitors (**Figure 1B**). In contrast, staining of NEUROD1, an early marker of the olfactory neuron lineage [31, 32], resulted negative (**Figure S1AB)**.

Despite the apparent cellular homogeneity of these cultures, we reasoned that bona fide neuronal progenitors of the HBC/GBC lineage might still be present at low proportions. To identify such cells, we performed scRNA-seq on the same eight samples (**Table S1**). After stringent quality control, we retained 146,462 cells for expression profiling, with an average detection of 5,232 genes per cell. To account for batch effects, including differences between donors, collection centers, and 10x chemistries, we integrated the eight samples using reciprocal PCA. Subsequent unsupervised clustering and visualization using uniform manifold approximation and projection (UMAP) indicated satisfactory removal of these confounders (**Figure S1C**). To increase sensitivity to rare cellular populations, we performed high-resolution clustering, using five neighbors to construct the graph, thereby allowing the formation of smaller clusters and yielding a total of 11 clusters (**Figure 1C**). We then estimated the cell-cycle phase using ccAFv2 and observed that cycling cells were arranged along a continuum from S phase to G1 (**Figure 1D**), indicating that, even after accounting for cell-cycle effects during normalization, cell-cycle state remained the main organizing factor in the UMAP.

To determine cellular identity, we integrated our dataset with a reference cellular taxonomy of the human OE. After integration with these reference datasets, we retained only label transfers with high confidence (see Methods). The majority of cells in our dataset aligned with populations annotated as fibroblasts/stromal cells (81%), with smaller fractions aligning with vascular smooth muscle cells (1,18%) and a minor subset aligning with pericytes (0.01%) and olfactory microvillar cells (0,19%) (**Figure 1E–G**). Cells with low-confidence predicted identities were classified as unknown (17,6%); however, these cells also most frequently aligned with fibroblasts in the reference dataset (**Figure 1E**). Notably, we did not identify any cellular cluster corresponding to the HBC/GBC/neuron axis associated with olfactory neurogenesis.

We next examined the expression of a manually curated list of molecular markers associated with the predicted cell populations, as well as markers linked to HBCs, GBCs, and immature and mature neurons (**Figure 2A,B**). All identified clusters showed high expression of genes associated with fibroblasts, including the type I collagen genes *COL1A1* and *COL1A2*, the proteoglycan decorin (*DCN*), and extracellular matrix components characteristic of fibroblasts/stroma such as fibulin-1 (*FBLN1*) and lumican (*LUM*). Other widely expressed genes included vimentin (*VIM*) and *PDGFRA*, both characteristic of mesenchymal cells. Additional genes detected across clusters included *PDGFRB* and *ACTA2*, markers typically associated with pericytes and vascular smooth muscle cells, respectively (**Figure 2A,B**). These genes are also known to be expressed in fibroblasts and may reflect transcriptional similarities between these lineages, potentially explaining the occasional classification of cells as pericytes or vascular smooth muscle cells in our previous analysis.

**Figure 2.**
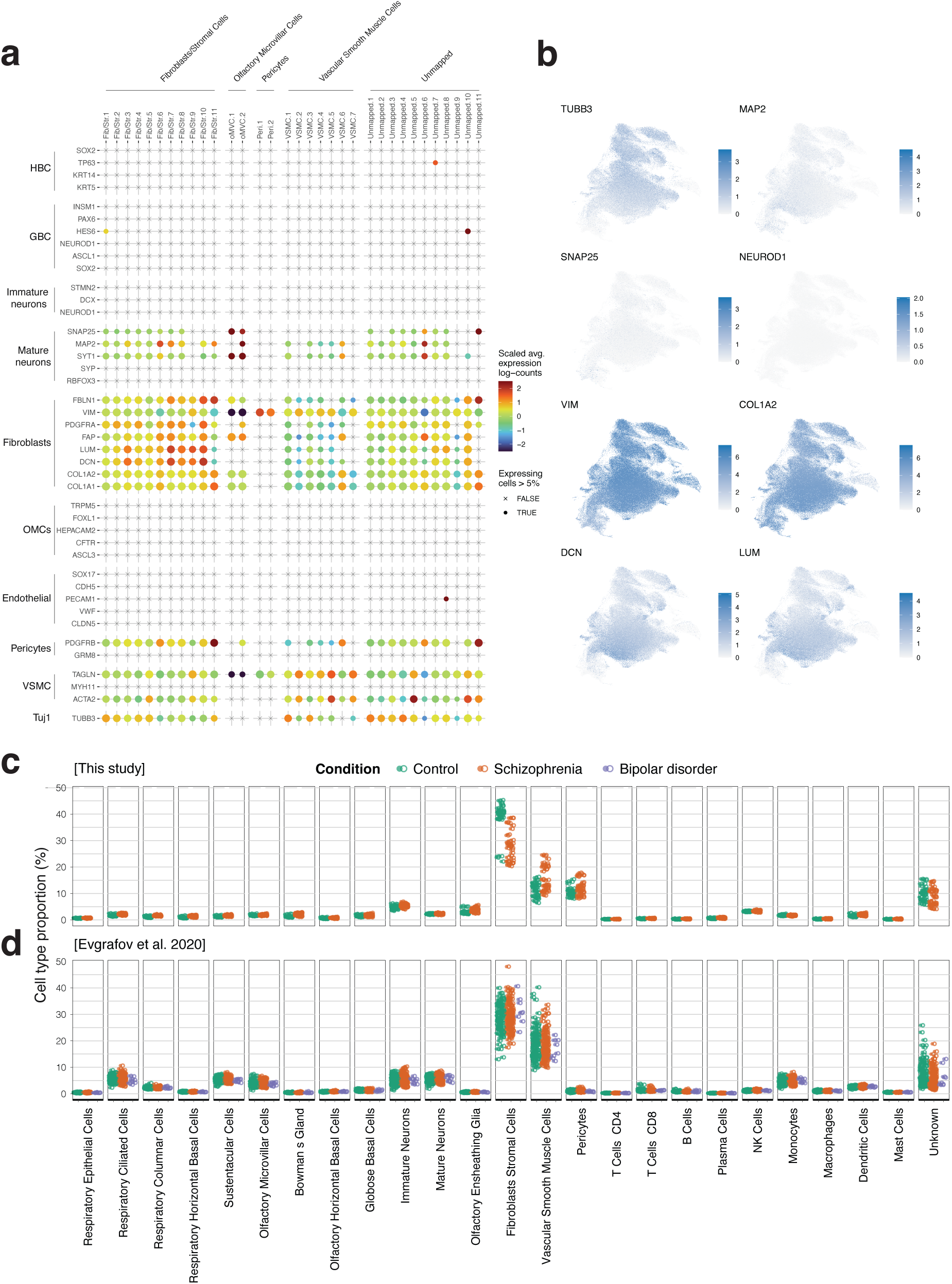
Marker gene expression and inferred cell-type composition in ON-derived cultures. (**a**) Dot plot showing the expression of selected marker genes across clusters identified in this study. Marker sets include OE lineages (horizontal basal cells, globose basal cells, immature and mature neurons), fibroblast/mesenchymal cells, olfactory microvillar cells, endothelial cells, pericytes, and vascular smooth muscle cells. Dot size indicates the fraction of expressing cells (>5%), and color represents scaled average expression. (**b**) UMAP feature plots showing the expression of representative neuronal (TUBB3, MAP2, SNAP25, NEUROD1) and mesenchymal (VIM, COL1A2, DCN, LUM) markers across the dataset. (**c**) Estimated cell-type proportions in this study based on deconvolution using a reference atlas of the human OE, shown for control and schizophrenia samples. (**d**) Cell-type proportions inferred from publicly available CNON-derived bulk RNA-seq data [22], including control, schizophrenia, and bipolar disorder samples.

Conversely, we did not detect expression of *SOX2*, a marker of basal progenitor and sustentacular cells [15], nor of canonical HBC markers *KRT5*, or *TP63*, nor of GBC markers including *PAX6*, *HES6*, *NEUROD1*, or *ASCL1*. Notably, *TUBB3* and other neuronal markers such as *MAP2* and *SNAP25* were detectable across clusters, albeit some of them at very low levels (**Figure 2A,B**). However, other canonical neuronal markers, including *RBFOX3*, *SYP*, *STMN2*, *DCX*, and *NEUROD1*, were not detected. While the previous analysis predicted a very small proportion of olfactory microvillar cells, canonical markers such as *ASCL3* were absent, providing limited support for their presence. Together, these results indicate that under these culture conditions the cells predominantly exhibit mesenchymal transcriptional identities, despite concomitant expression of a small number of genes commonly associated with neurons. In summary, both immunostaining and single-cell transcriptomic profiling indicate that these adherent cultures of nasal exfoliates are largely composed of cells of mesenchymal identity.

To assess the generalizability of our findings, we generated bulk RNA-seq data from passage 3 cultured nasal exfoliates derived from 10 neurotypical individuals and 10 schizophrenia patients. We then applied Scaden, a deep learning–based deconvolution method, using single-cell RNA-seq data from the human OE as reference [31], to estimate cell-type proportions. Consistent with our single-cell RNA-seq analysis, all samples showed a predominant contribution of fibroblast/mesenchymal cells, followed by vascular smooth muscle cells and pericytes. The method estimated only minor contributions (<5%) from additional cell types in the bulk profiles (**Figure 2C**).

We next analyzed publicly available bulk RNA-seq data from cell cultures derived from nasal biopsies of 118 control individuals, 144 schizophrenia patients, and 8 bipolar disorder patients [22]. This system, which involves growth in Matrigel using medium 4506, has been reported to generate neuron-like cells, although lacking markers of mature olfactory neurons [39]. Application of the same deconvolution framework yielded a highly similar cellular composition, again characterized by a predominant fibroblast/mesenchymal signature. Together, these results indicate that mesenchymal lineages consistently dominate primary cultures derived from nasal exfoliates across protocols and cohorts (**Figure 2D**).

### The potential of ON-derived cultures to model brain diseases

One approach to evaluate the suitability of a model system for studying human diseases is to quantify the expression of disease-associated genes within the relevant cell types. Intrinsic molecular alterations can only be captured if the implicated genes are expressed in the model. To assess the potential of the different cell types present in the OE to detect such alterations, we analyzed the expression patterns of curated disease-associated gene sets across cell populations in the in vivo human OE [31]. We studied 25 gene sets from Mato-Blanco et al. 2025 [7], considering a wide spectrum of brain diseases, including malformations of cortical development (such as microcephaly, lissencephaly, heterotopia and polimicrogyria) and neurodevelopmental disorders (such as schizophrenia, bipolar disorder, autism spectrum disorder, and major depression).

First, to obtain a detailed view of the enrichment patterns of each of the 25 disease-associated gene sets across OE cell types, we applied the AUCell algorithm [40]. Consistent with previous findings in the developing human and macaque brain [7, 41], genes associated with neurodevelopmental disorders were enriched in neuronal populations, with autism spectrum disorder–associated genes showing preferential enrichment in immature rather than mature neurons. In contrast, genes linked to Alzheimer’s disease showed the expected enrichment in immune cell signatures. Gene sets associated with malformations of cortical development displayed more heterogeneous patterns: while polymicrogyria and lissencephaly were enriched in immature neuronal populations, other conditions such as developmental dyslexia, hydrocephalus, and cobblestone lissencephaly showed enrichment in olfactory ensheathing glial cells. Notably, genes associated with microcephaly, polymicrogyria, and rare malformations of cortical development also showed enrichment in GBCs, paralleling previously reported enrichments in cortical intermediate progenitor cells [7] and supporting a transcriptomic similarity between these progenitor populations across distinct neurogenic niches. In contrast, mesenchymal cell populations were largely depleted of enrichment signals across this comprehensive set of brain disease–associated gene sets.

Despite the overall enrichment of disease-associated genes in neuronal and neurogenic populations, individual genes may still be expressed across multiple cell types. To quantify this, we measured the proportion of disease-associated genes expressed across increasing fractions of cells within each cell type. As expected, globose basal cells, as well as immature and mature neurons, showed the highest proportions of expressed disease-associated genes, even at the lowest detection threshold (≥5% of cells), below which expression was considered noise (see Methods). We summarized this behavior by calculating the area under the curve (AUC) of the cumulative expression fractions. Consistent trends were observed across individual disease categories (**Figure 3B**). In contrast, fibroblast/mesenchymal cells consistently exhibited lower proportions of expressed disease-associated genes (**Figure 3B,C**), particularly at intermediate expression thresholds. At higher thresholds, however, differences between cell types became less pronounced, reflecting the inherent sparsity of single-cell RNA-seq data, where few genes are detected in a large fraction of cells. Together, these results indicate that disease-associated gene expression is preferentially enriched and more consistently detected within neurogenic lineages, while mesenchymal populations contribute comparatively limited signals.

**Figure 3.**
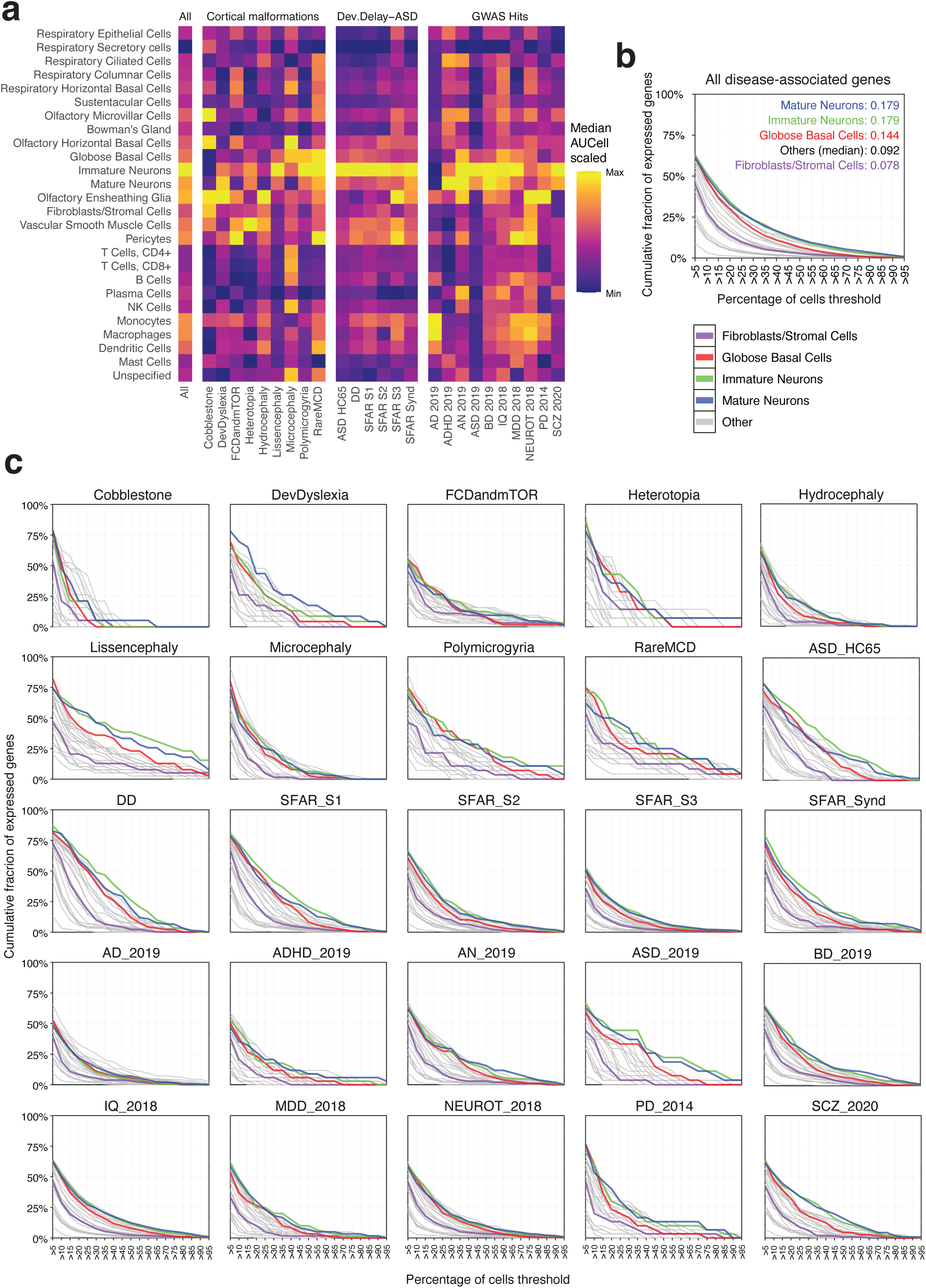
Expression patterns of disease-associated gene sets across OE cell types. (**a**) Heatmap showing AUCell enrichment scores for disease-associated gene sets across cell types of the human OE. Gene sets are grouped into cortical malformations, neurodevelopmental disorders (Dev.Delay–ASD), and GWAS-derived traits. Values represent scaled median AUCell scores per cell type. (**b**) Cumulative fraction of genes from all disease-associated gene sets expressed above increasing cell-percentage thresholds (≥5% to ≥95%) across major cell types. Lines correspond to fibroblasts/stromal cells, globose basal cells, immature neurons, mature neurons, and other cell types. Area under the curve (AUC) values are indicated for each cell type. (**c**) Cumulative fraction of expressed genes for individual disease-associated gene sets across increasing cell-percentage thresholds. Each panel corresponds to a specific disease or trait, including cortical malformations, neurodevelopmental disorders, and GWAS-derived gene sets. Lines represent different cell types as in (**b**).

## Discussion

Neurogenesis in the OE provides a unique opportunity to investigate neurodevelopmental processes across the lifespan, an advantage long recognized in the field. Sensory neurogenesis has traditionally been assumed to originate from horizontal and globose basal cells; however, our results indicate that, under commonly used experimental conditions, the cellular populations interrogated are frequently dominated by mesenchymal lineages instead.

One important source of variability across studies lies in both the anatomical location of the biopsy and the explant culture methodology. Feron et al. showed that the likelihood of detecting active olfactory neurogenesis in healthy individuals increases when biopsies are restricted to the dorsoposterior nasal septum and the opposing surface of the superior turbinate, and when the epithelium is cultured as thin slices rather than larger tissue fragments [42]. However, although the OE is most consistently enriched in the superior turbinate, this region is difficult to access in routine procedures and is typically sampled only during surgery under general anesthesia [39, 43]. Consequently, many studies rely on more accessible regions, such as the anterior and medial portions of the middle turbinate, where neuronal populations can be detected but at lower frequency [35, 42, 44]. Differences in sampling location may therefore lead to substantial variability in the abundance of neuronal progenitors across studies and individuals.

However, our results indicate that this source of variability alone cannot account for the observed cellular composition. Across multiple participants, with samples obtained by different operators and processed in independent centers, we consistently observed a strong bias toward mesenchymal cell populations. Together with previous reports, this points to an intrinsic effect of in vitro culture conditions, which appear to favor the rapid expansion and early dominance of fibroblast-like cells, likely originating from the lamina propria, a connective tissue-rich compartment adjacent to the olfactory epithelium. A similar drift toward mesenchymal progenitors has been observed in neurosphere-based cultures, suggesting that this bias reflects general properties of current culture paradigms. Notably, the potential of mesenchymal stem cells from olfactory and non-olfactory nasal lamina propria to form spheres and differentiate into both osteogenic and neuronal lineages, has been observed [45, 46]. Stem cells derived from this compartment share membrane markers with bone marrow–derived stem cells and have therefore been termed olfactory ectomesenchymal stem cells [45, 47].

Importantly, our data indicate that commonly used neuronal markers may be insufficient to resolve this issue. βIII-tubulin immunoreactivity has often been interpreted as evidence of neuronal identity [10, 16, 18, 19, 39]; however, in the absence of orthogonal markers and cell-type–resolved transcriptomic validation, its expression alone does not reliably indicate bona fide neuronal differentiation or GBC–derived neurogenesis. In our dataset, a subset of neuronal markers can be co-expressed alongside a broad repertoire of mesenchymal genes, underscoring the risk of misclassification in serum-expanded cultures, particularly at later passages.

Compared to the GBC-to-neuron axis, mesenchymal cells offer more limited potential to reveal or investigate neurodevelopmental molecular alterations associated with brain disorders [48]. Nevertheless, this tissue still present advantages to reveal more systemic molecular alterations associated with human brain disorders. As an accessible source of highly proliferative and relatively homogeneous cells, they provide a tractable model in which to examine biological processes that may partially overlap with those operating in neural progenitors. In this context, previous studies have reported alterations in pathways related to WNT5A signaling, mitosis, and cell-cycle regulation in schizophrenia, as well as oxidative stress-related abnormalities across schizophrenia, Alzheimer’s disease, and Parkinson’s disease [16, 17, 22, 42, 49]. Additional work has described differences in cell morphology, cytoskeletal organization, synaptic organization, and extracellular matrix interactions in schizophrenia and bipolar disorder [14, 18, 19, 50, 51]. Likewise, properties of specific neurotransmitter systems, including cannabinoid and serotonin receptors, have been investigated in these cultures and linked to neuropsychiatric phenotypes and antipsychotic treatment response in schizophrenia [20].

In contrast to in vitro systems, recent work using single-nuclei RNA-seq directly on frozen olfactory epithelium swabs has demonstrated preservation of cellular diversity, including neuronal progenitors and mature neuronal populations, in both humans and pigs [15]. These findings support a shift toward direct profiling approaches to capture the native cellular landscape, alongside the development of improved strategies to isolate and culture bona fide neuronal progenitor populations. Notably, the neurogenic axis of the OE mirrors patterns of disease gene enrichment reported in the developing cortex, with progenitor and neuronal populations preferentially expressing disease-associated genes. This parallel further supports the use of the OE as an accessible system to interrogate neurodevelopmental risk. Together, such efforts will be essential in order to properly exploit the great potential that the OE holds for the study of neurodevelopmental disorders.

## Funding for open access charge

Instituto de Salud Carlos III [CP20/00064], with co-financing by European Funds for the Miguel Servet Contract.

## Authoŕs contributions

X.M-B., performed the bioinformatic analyses for this study. S.B., AC and E.V performed all experimental work. G.M. A.T., A.R., V.S-G, and A.M recruited patients. Z.P, M.B-C, MJA, MG and CS-A performed nasal biopsies. P.R. coordinated samples adquisition and explant culturing at Hospital del Mar. G.M. coordinated samples adquisition and explant culturing at Hospital Universitari Institut Pere Mata. G.S. and P.R. conceived the study and wrote the manuscript. G.S. and P.R. supervised the study. All authors read and approved the manuscript.

## Data and materials availability

All data needed to evaluate the conclusions in the paper are present in the paper and/or the Supplementary Materials. All code to reproduce the analyses presented in the paper will be available at GitHub uppon publication.

## Methods

### ON sampling and culture

Four participants were recruited from the Hospital Universitari Institut Pere Mata (HUIPM) and four from Hospital del Mar in Spain. Patients were recruited from the outpatient psychiatric units at HUIPM, and HC subjects were patient’s friends, nongenetic relatives and university students directly interviewed by an experienced psychiatrist to exclude subjects with a past or current history of psychiatric disorders. Diagnoses of schizophrenia were established in accordance with the Diagnostic and Statistical Manual of Mental Disorders, Fourth Edition (DSM-IV) criteria. Ethical approval was granted by the hospital’s ethics review committee, and all participants provided written informed consent (Ref:2022/10297/I and 037/2022, respectively). After humidification of the nasal cavity, two separate sterile interdental brushes were used to collect samples from the lower and middle turbinates. Each brush was immediately placed into an Eppendorf tube containing 250 μl of cold Dulbecco’s Modified Eagle Medium/Ham’s F-12 (DMEM/F12; GibcoBRL) supplemented with 10% fetal bovine serum (FBS), 2% glutamine, 1% anti-anti, and 1% penicillin–streptomycin. This procedure was performed independently for each nostril.

Nasal exfoliates were maintained in cold supplemented DMEM/F12 and mechanically dissociated to obtain single-cell suspensions. Primary cultures were established and maintained for 3 weeks in supplemented medium at 37 °C in a humidified incubator with 5% CO₂ before passaging into culture flasks (Thermo Scientific, Madrid, Spain).

Cells were dissociated using 0.25% trypsin (GibcoBRL), replated at a density of 4,000 cells/cm² into 75 cm² flasks, and cultured in supplemented medium. Cultures were subsequently expanded through serial passaging. After harvest, cells were cryopreserved in aliquots using DMEM/F12 containing 2% glutamine, 1% penicillin–streptomycin, 20% FBS, and 10% dimethyl sulfoxide (DMSO; Sigma-Aldrich, Madrid, Spain), and stored in liquid nitrogen.

Frozen aliquots of OE cells at passage 3 were used as the starting material for all downstream experiments, including immunohistochemistry, bulk and single-cell RNA-seq.

### Immunohistochemistry

Cells were seeded at 1 × 10⁵ cells ml⁻¹ on sterilized glass coverslips. At ∼70% confluency, cells were washed with phosphate-buffered saline (PBS) and fixed with 4% paraformaldehyde (PFA) for 15 min at room temperature. Cells were then permeabilized with 0.2% Triton X-100 for 10 min and blocked in PBS containing 2% bovine serum albumin (BSA), 2% donkey serum (DS), and 0.1% Triton X-100 for 1 h. Primary antibodies were diluted in blocking buffer and incubated overnight at 4 °C (monoclonal mouse anti-TUBB3, Invitrogen (TU-20) 1:1000; monoclonal mouse anti-Ki-67 DAKO (M7240), 1:500; polyclonal rabbit anti-PDGFRA, Thermo (710169) 1:100), and anti-NEUROD1 Proteintech (12081-1-AP) 1:50. Coverslips were washed three times with PBS and incubated with Alexa Fluor 568–conjugated secondary antibodies for 2 h at room temperature in the dark (anti mouse Abcam (ab175473); anti rabbit Abcam (ab150077)). Nuclei were stained with DAPI and coverslips mounted using Fluoaqueous mounting medium. Images were acquired with a Leica SP8 confocal laser scanning microscope using 20× and 40× objectives.

### sc-RNA sequencing

Adherent cultures (80–90% confluency) were dissociated to generate single-cell suspensions for single-cell RNA sequencing. Cells were washed twice with PBS w/o Ca²⁺ and Mg²⁺ and incubated with pre-warmed 1× trypsin-EDTA at 37 °C for ∼8 min until detachment was observed. Trypsinization was quenched with four volumes of complete growth medium, and cells were collected and centrifuged at 300g for 3 min. The cell pellet was washed twice with PBS supplemented with 0.1% bovine serum albumin (BSA), filtered through a 37 µm cell strainer, and resuspended in PBS containing 0.1% BSA. Cell concentration and viability were determined using trypan blue staining and an automated cell counter, and samples were adjusted to the required concentration and maintained on ice prior to processing.

Single-cell suspensions were prepared as described above and delivered to the institutional Genomics Facility. Cells were resuspended in PBS supplemented with 0.1% BSA at a concentration of 1600 cells μl⁻¹, following the facility’s standard submission guidelines. Approximately 20,000 cells were targeted for recovery during library preparation. Single-cell libraries were generated using the 10x Genomics Next GEM Single Cell 3′ Reagent Kits v3.1 according to the manufacturer’s protocol and the standard workflow of the Genomics Facility. Libraries were subsequently sequenced following the recommended 10x Genomics parameters.

### Single-cell RNA-seq processing, quality control, and integrated analysis

Demultiplexed FASTQ files were processed independently for each of the eight single-cell libraries using Cell Ranger (v10.0.0; 10x Genomics, 2024) with the GRCh38-2024-A human reference transcriptome. Ambient RNA contamination was subsequently removed independently for each sample using CellBender (v0.3.2; [52]) applied to the raw count matrices. CellBender runs were evaluated using the standard log and diagnostic outputs, and libraries showing unstable optimization or implausible droplet classification were re-run with adjusted learning rate, number of epochs, and, when necessary, manually specified droplet priors. For downstream analyses, the corrected matrix corresponding to false positive rate 0.01 was retained for each sample.

Accepted CellBender-corrected matrices were imported into Scanpy (v1.12; [53]), and standard quality-control metrics were computed, including mitochondrial, ribosomal, and hemoglobin-associated counts. Cells were filtered using minimum thresholds of 1,000 total counts and 100 detected genes, and genes with no detected expression after cell filtering were removed. Doublet scores were computed using Scrublet (v0.2.3; [54]) and scDblFinder (v1.20.2; [55]) as part of quality-control assessment. Cell-cycle state was inferred with ccAFv2 (v2.0.6; [56]), and a permissive cell-cycle class assignment (threshold = 0) was carried forward for downstream integrated analysis.

The eight processed samples were then combined into a single dataset for integrated analysis. The final analysis used Seurat (v5.3.0), using SCTransform normalization (v0.4.2; [57]) and RPCA-based integration within the Seurat framework [58]. Sample identity was used as the integration variable, whereas the cell-cycle class was included in vars.to.regress during SCTransform. UMAP visualization and graph-based clustering were performed on the final integrated representation using a 5-nearest-neighbor graph and Leiden clustering at resolution 0.2 [59]. Cluster interpretation was guided by the expression of canonical markers representing olfactory epithelial, neuronal, mesenchymal/stromal, vascular, and immune populations.

### Reference-based cell type annotation

Cell identities were further evaluated by mapping the integrated query dataset onto an external annotated human OE reference from Durante et al. (2020) [31]. The reference dataset was filtered to exclude cells labeled as Unspecified and was further restricted to Patient 2 for the mapping analysis used in the manuscript. The query dataset was preprocessed in Seurat by splitting the RNA assay by sample and applying SCTransform, whereas the reference dataset was SCTransformed independently, followed by PCA, construction of a neighbor graph using dimensions 1–30, and UMAP embedding using the same dimensions.

Label transfer was performed using FindTransferAnchors and TransferData with `normalization.method = “SCT”`, `dims = 1:30`, and `reference.reduction = “pca”`, transferring the reference CellType labels to the query cells. To project query cells into the reference manifold, a reference UMAP model was computed with `return.model = TRUE`, after which the query was projected using IntegrateEmbeddings and ProjectUMAP. For downstream interpretation, cells with maximum transfer score < 0.8 were classified as Unmapped, whereas cells above this threshold retained their transferred identity.

### Bulk RNA-seq

Adherent cultures (80–90% confluency) were dissociated to generate cell suspensions. Cells were washed twice with PBS w/o Ca²⁺ and Mg²⁺ and incubated with pre-warmed 1× trypsin–EDTA at 37 °C for ∼8 min until detachment. Trypsinization was quenched with complete growth medium, and cells were collected by centrifugation at 300g for 3 min. The cell pellet was washed twice with PBS. Total RNA was extracted using the RNAqueous Total RNA Isolation Kit (Thermo Fisher Scientific) according to the manufacturer’s instructions. Briefly, cell pellets were lysed, and RNA was purified using a fiber filter-based purification system, followed by washing and elution of total RNA. Fifty-two RNA samples were submitted to HaploX GeneTech (Hong Kong) for library construction and sequencing. Libraries were sequenced on an Illumina NovaSeq X Plus platform with paired-end 150 bp reads, generating approximately 9 Gb of sequencing data per sample.

### Bulk RNA-seq processing and gene-level quantification

Raw paired-end bulk RNA-seq reads were first assessed using FastQC (v0.11.5), and quality reports were aggregated with MultiQC [60]. A genome index was generated with STAR (v2.7.9a; [61]) from the GRCh38 human reference genome and Ensembl GRCh38 gene annotation (release 104). Reads were aligned with STAR, and alignment files were converted from SAM to BAM format using SAMtools (v0.1.19; [62]). Read-group information was then added with Picard (v2.18.12), followed by BAM sorting with SAMtools and duplicate removal using Picard MarkDuplicates. Gene-level read counts were obtained with featureCounts from the Subread package (v1.6.4; [63]) using exon-based assignment and summarization at the gene_id level. Sample-specific count files were merged into a single count matrix, and genes with no expression across all samples were removed prior to downstream analyses.

### Bulk RNA-seq deconvolution

Cell-type deconvolution of bulk RNA-seq profiles was performed with Scaden (v1.1.2; [64]) using a human OE single-cell reference derived from Durante et al. (2020) [31]. Two bulk datasets were analyzed: the in-house bulk RNA-seq cohort generated from cultured nasal exfoliates (using reads from each sequencing lane as pseudo-replicates) and an external bulk RNA-seq dataset from cultured OE-derived cells (CNON) reported by Evgrafov et al. (2020) [22]. For Scaden input preparation, bulk and single-cell datasets were harmonized on the intersection of shared genes. Single-cell reference profiles were normalized on a per-cell basis before simulation of pseudo-bulk mixtures. The Scaden training configuration used 2,000 cells per simulated sample, 5,000 simulated samples, var_cutoff = 0.1, batch_size = 128, learning_rate = 1×10-4, 5,000 training steps, and seed = 1997. The cell annotation label “Unspecified” was used as the unknown class. Final predicted cell-type proportions were obtained from the Scaden output tables generated for each bulk dataset and used for downstream visualization and comparison across samples.

### Disease-associated gene expression and enrichment in the primary OE

Disease-associated genes were curated as described elsewhere in Mato-Blanco et al. (2025) [7] and organized into three major groups, Cortical Malformations, Dev.Delay-ASD, and GWAS Hits, together with a combined All category. These gene sets were analyzed in the human OE reference dataset from Durante et al. (2020) [31]. Raw counts were retained for prevalence analyses, and the data were normalized to a total of 10,000 counts per cell and log-transformed for summary analyses. Disease-associated genes were intersected with the genes present in the reference dataset before downstream quantification.

To quantify how broadly disease-associated genes were represented across reference cell types, expression statistics were computed separately for each CellType and gene, including the fraction of cells with detectable expression. Genes were then binned according to the percentage of expressing cells in 5% increments from 0 to >95%, and cumulative fractions were calculated across thresholds for each disease category and cell type. These cumulative distributions were used for summary comparisons across cell populations.

In parallel, disease-associated gene sets were scored directly in the Durante reference using AUCell (v1.28.0), as originally described within the SCENIC framework [40]. AUCell scores were then summarized by CellType, using scaled median values as the principal cell-type-level summary of disease-gene enrichment.

---

**Supplementary Figure 1.**
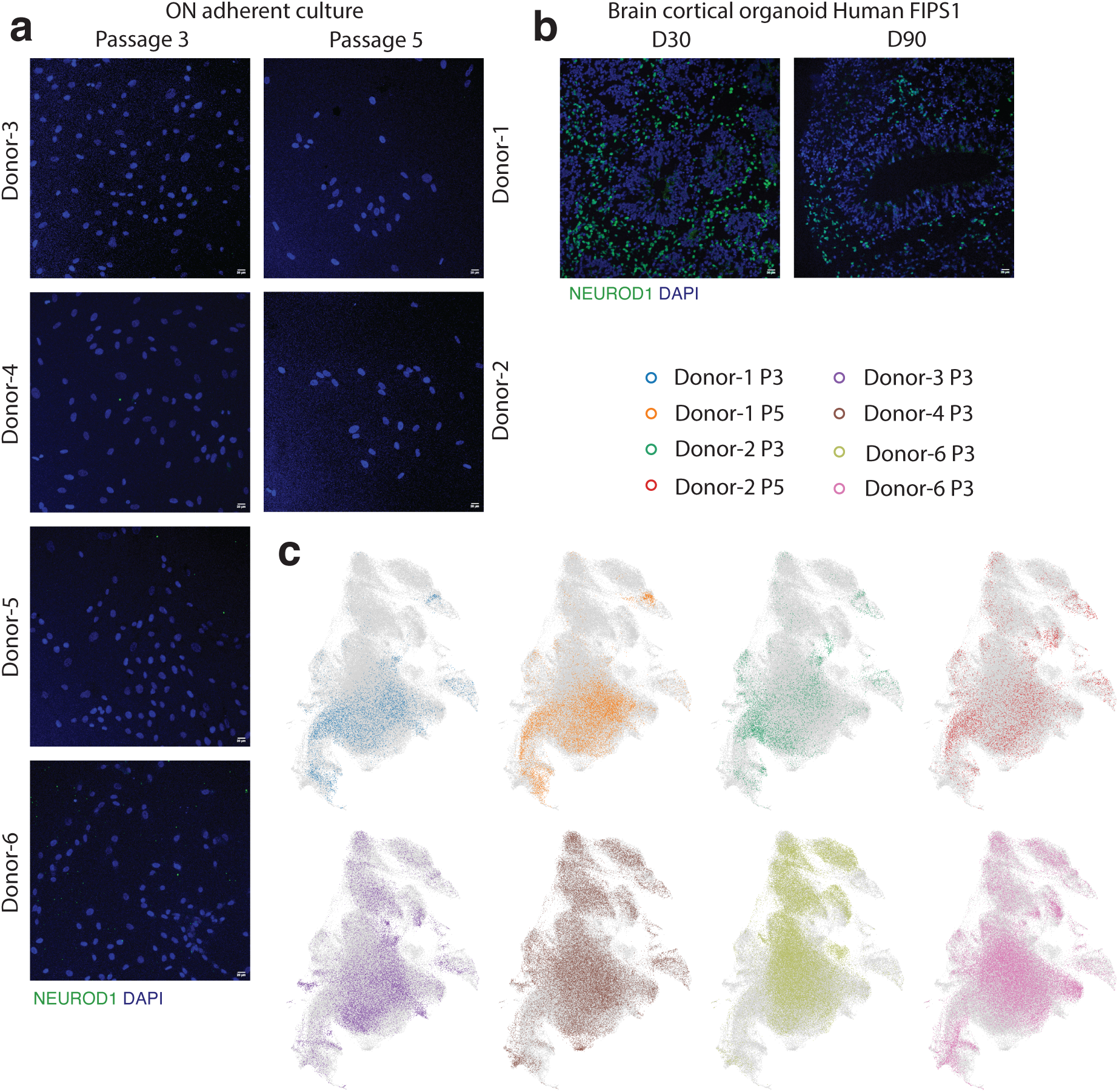
(a) Immunofluorescence staining for NEUROD1 (green) and DAPI (blue) in adherent cultures derived from nasal exfoliates at passage 3 and passage 5. **(b)** Immunofluorescence staining for NEUROD1 (green) and DAPI (blue) in human cortical organoids (FIPS1 line) at day 30 (D30) and day 90 (D90), used here as a positive control for antibody specificity. **(c)** UMAP visualization of single-cell RNA-seq data colored by sample identity, including control and schizophrenia samples across passages.

**Table S1.** Description of samples.

| Table S1. Description of samples |  |  |  |  |
| --- | --- | --- | --- | --- |
| Donnor | Diagnose | Passage | Center | Operator |
| 1 | CTRL | 3 | HMRI | M.B-C |
| 1 | CTRL | 5 | HMRI | M.B-C |
| 2 | CTRL | 3 | HMRI | M.B-C |
| 2 | CTRL | 5 | HMRI | M.B-C |
| 3 | CTRL | 3 | HMRI | Z.P. |
| 4 | CTRL | 3 | HUIPM | MJA/MG/CS-A |
| 5 | SCZ | 3 | HMRI | Z.P. |
| 6 | SCZ | 3 | HUIPM | MJA/MG/CS-A |

